# Inter- and Intra-individual Variability in Oral Food Processing and Its Impact on Aroma Release

**DOI:** 10.64898/2026.05.05.721895

**Authors:** I. Andriot, D. Grossiord, N. Béno, T. Chabin, H. Labouré, G. Lucchi, C. Martin, O. Mourabit, J.A. Piornos, L. Saint-Georges, C. Salles, I.C. Trelea, C Peltier

**Affiliations:** Université Bourgogne Europe, Institut Agro, CNRS, INRAE, UMR CSGA, 21000 Dijon, France; Probe Research Infrastructure, ChemoSens facility, CNRS, INRAE, 21000 Dijon, France; Université Paris-Saclay, INRAE, AgroParisTech, UMR SayFood, F-91120, Palaiseau, France

## Abstract

Aroma perception during food consumption results from the combined effects of food composition, oral processing (such as chewing and saliva action), the release and transport of volatile compounds toward the olfactory epithelium, followed by cognitive integration in the brain. Recent advances in real-time analytical techniques, particularly Proton Transfer Reaction-Time-of-Flight Mass Spectrometry (PTR-ToF-MS), enable *in vivo* monitoring of aroma release with high temporal resolution and have become widely used for analyzing the composition of exhaled air. However, the interpretation of aroma release kinetics remains challenging due to substantial intra- and inter-individual variability caused by differences in physiology, anatomy, oral behavior, and respiratory patterns.

In this context, the present study was designed to quantify aroma release associated with different food oral processing (FOP) mechanisms, such as chewing and swallowing, using simple model matrices containing a single aroma compound, and to document inter- and intra-individual variability among subjects.

Real-time PTR-MS measurements were combined with self-reported oral events and simultaneous respiratory monitoring to analyze aroma release from aqueous solutions and gummy discs flavored with isoamyl acetate. The results showed that inter-individual variability was higher than intra-individual variability and allowed its quantification in aroma release. Significant differences in aroma release kinetics were observed depending on FOP protocols. The importance of considering swallowing events when analyzing aroma release data was also highlighted.

## I. Introduction

Aroma release during food consumption results from a dynamic interplay between food components, oral processing, and the transport of volatile compounds to the olfactory epithelium in the nasal cavity. Recent advances in real-time analytical techniques—most notably Atmospheric Pressure Chemical Ionization-Mass Spectrometry (APCI-MS) and Proton Transfer Reaction-Mass Spectrometry (PTR-MS) —have enabled *in vivo* monitoring of aroma release with high temporal resolution. PTR-MS, particularly when coupled with a Time-of-Flight analyzer (PTR-ToF-MS), is highly suited for nosespace analysis thanks to its sensitivity and rapid response time (Jordan et al., 2009; Le Quéré and Lucchi, 2022). This approach has been widely applied to characterize volatile organic compounds (VOCs) released during the consumption of diverse food matrices, including alcoholic solutions, wine, coffee, apples, model cheeses, and chocolate (Acierno et al., 2019; Andriot et al., 2024; Arvisenet et al., 2019; Charles et al., 2015; Deleris et al., 2011; Deuscher et al., 2019; Heenan et al., 2012; Romano et al., 2014; Ting et al., 2016). A recent review (Le Quéré and Schoumacker, 2023) emphasized that release kinetics strongly depend on the food matrix and oral processing mechanisms. Importantly, substantial inter-individual variability in aroma release complicates the interpretation of averaged data. This variability arises from differences in physiology and oral behavior, such as mastication patterns, saliva incorporation and swallowing (Burdach and Doty, 1987), the latter often triggering sharp increases in VOC flow through the nasal cavity. Anatomical differences (oral, nasal and pharyngeal cavity volumes), physiological parameters (salivary flow rate, saliva composition), and aroma–mucosa interactions governed by air/mucosa partition coefficients further contribute to these differences (Déléris et al., 2016; Pedrotti et al., 2019).

Although absolute signal intensity may vary across repetitions, the temporal pattern of release curves tends to remain homogeneous within individuals after normalization (Deuscher et al., 2019; Repoux et al., 2012; Romano et al., 2014). However, respiratory cycles and swallows introduce significant complexity into PTR-MS signals leading to irregularities and difficulties to predict the curves. To manage this variability, most studies simplify aroma release curves into a small number of summary measures, such as maximum intensity (Imax) or area under the curve (AUC). While these parameters are useful, they lose the rich temporal information contained in the full curve. Other approaches rely on smoothing, temporal averaging, or binning procedures, such as maximal envelopes (Doyennette et al., 2014) or breathing-cycle detection (Peltier et al., 2024). Finally, mechanistic models aim to reproduce these signals using differential equations derived from physics (such as diffusion or convection equations), while accounting for breathing and the oral processing of food. These models were validated by averaging out inter-individual and FOP-related variability using datasets obtained with complex foods (Doyennette et al., 2014).

In this context, the present study was designed to quantify aroma release associated with different food oral processing (FOP) mechanisms, such as chewing and swallowing, using different simple model matrices containing a single aroma compound. Therefore, future mechanistic modeling of individual aroma release kinetics could be supported by incorporating the effects of FOP mechanisms. In addition, the study sought to characterize and identify the sources of intra- and inter- individual variability in aroma release, in order to enable model generalization across the overall population. Based on the literature, we hypothezised that swallowing events induce sharp increases in aroma release due to increased retronasal airflow, and that chewing increases aroma release through mechanical disruption of the matrix and facilitates VOC transfer from the oral to the nasal cavity.

This paper presents the analysis of the obtained dataset, focusing on:

i. the experimental impact of different FOP mechanisms on aroma release for various matrices;
ii. the characterization of intra- and inter-individual variability in aroma release

## 2. Material and methods

### 2.1. Experimentation

#### 2.1.1. Ethics

This study was approved by CERUBFC-2025-02-20-017 (Comité d’Éthique de la Recherche de l’Université Bourgogne Franche-Comté). All procedures complied with the Declaration of Helsinki, and written informed consent was obtained from all subjects before the beginning of the experiments.

#### 2.1.2. Subjects

Recruitment was conducted by email through the PanelSens database of the CSGA (Center of Taste and Feeding Behavior), which includes contact information for volunteers interested in participating in studies. To participate in this study, it was mandatory not to have any swallowing or Ear, Nose, and Throat (ENT) disorders or not be taking any medication or not having any medical conditions that could alter sensory perception or physiological functions related to the study.

The study started on June 2^nd^ 2025 and ran until August 8^th^ 2025. It consisted of three one-hour tasting sessions per subject, spread over one to two months depending on subject availability. Subjects were compensated with a 10€ voucher per 1-hour session. The recruited panel was finally composed of 30 subjects (15 men, 15 women) aged from 22 to 57 years old.

#### 2.1.3. Products

Two different products were tasted in this study: a water solution containing isoamyl acetate and a disc gummy containing the same aroma compound.

First, an isoamyl acetate solution (0.4 g.L^-1^) was prepared by dissolving 40 mg of isoamyl acetate ( ≥ 97%, Food Grade, FFC Food Flavouring Compound, Sigma-Aldrich, France) in 80 mL of Evian® water (Danone, France) in a closed amber flask and stirring for 1 hour at room temperature (∼21°C). After that, the solution was transferred in a 100 mL volumetric flask and the volume was adjusted with Evian® water to 100 mL. The solution of isoamyl acetate (0.4 g.L^-1^) was prepared each day before the sessions. This solution was presented to the panelists either in cups (“solution” samples) or by a gustometer (“gusto” samples).

Second, a gelatin gummy disc (diameter 23 mm, thickness 0.4 mm) containing 10% w/w pork gelatin (gelatin in powder, Vahiné, France) and isoamyl acetate (0.4 g.L^-1^) was prepared. An isoamyl acetate solution (2 g.L^-1^) was prepared by dissolving 200 mg of isoamyl acetate in 80 mL of Evian® water (Danone, France) in a closed amber flask and stirring for 2 hours at room temperature (∼21°C). After that, the solution was transferred in a 100 mL volumetric flask and the volume was adjusted with Evian® water to 100 mL. The gelatin powder (45 g) was first hydrated in 200 g of Evian® water under stirring for 5 minutes in a Thermomix cooking robot at room temperature. Then, 150 g of warm Evian water were added and the solution was heated to 37°C. Once this temperature reached, 100 g of the solution containing isoamyl acetate (2 g.L^-1^) was added and the mixture was stirred for 2 minutes. The mixture was transferred to silicone disc-shaped molds (2 mL in each well), covered with cling film and stored overnight at 4°C. The gummy discs were put in small cups covered with caps (one gummy disc per cup), and were taken out the cold chamber 15 minutes before each sensory session. Each batch of gummy discs were kept in the cold chamber for up to three days, after which they were disposed if not consumed.

#### 2.1.4. Tasting procedure

The experiment combined real-time PTR-MS nosespace measurements during the tasting of aqueous solutions and gummy discs flavored with isoamyl acetate, with either imposed or self-reported oral events (Figure 1a). This aimed to quantify the impact of different FOP mechanisms on aroma release.

**Figure 1:**
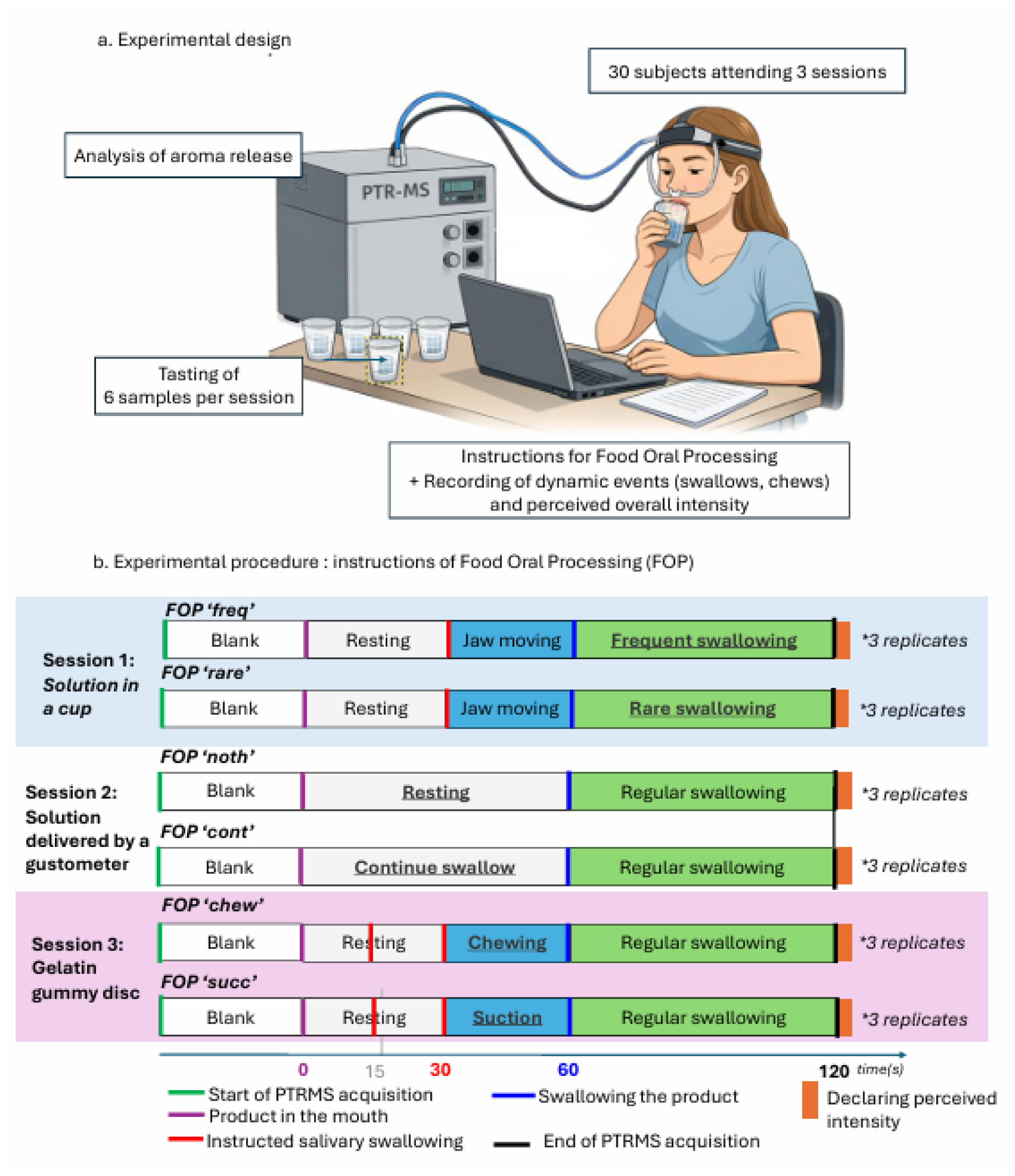
Experimental design (a) and procedure with instructions (b)

In this study, the term ‘delivery mode’ refers to the way in which the product was administered, regardless of its physical form. Although only two products were tested (a solution and a gel), three delivery modes were considered. Each session was dedicated to the tasting with a single delivery mode repeated six times: during the first session, the solution presented in a cup (“solution in a cup”) was evaluated; the solution delivered by the gustometer was evaluated in the second session (“solution delivered by gustometer”); and the gelatin gummy disc in the third session (“gelatin gummy disc”). The order of presentation of delivery modes (“solution in a cup”, “solution delivered by gustometer” or “gelatin gummy disc”) was fixed for organizational reasons.

First, the subjects were connected to PTR-ToF-MS via two canulae that were mounted on a helmet. The tip of these canulae (a few millimeters) were put in the participant’s nostrils so the nosespace could be aspirated and a signal was observed on the PTR-ToF-MS. The participants tasted a warm-up sample consisting of the delivery mode used in the session (solution in a cup, solution delivered with gustometer or gummy disc). Then, they tasted the sample 6 times and the concentration of aroma compounds was monitored during 120 s (Figure 1b).

For each tasting, subjects first breathed normally for 30 s to record a baseline signal prior to product consumption (−30-0s interval), then followed a predefined protocol to consume the product (Figure 1b). After the 30 s corresponding to the baseline signal, subjects placed the product in their mouth, and pressed ENTER on the keyboard to play a sound, enabling the experimenter to synchronize PTR-ToF-MS measurements with recorded events such as swallows or chews. The subjects then followed the instructions on the screen to complete a standardized 120 seconds tasting protocol detailed in the next subsection. In addition to the imposed FOP experiments, subjects declared any accidental chews and/or swallows that were not included in the protocol during the tasting, by pressing keys (‘m’ for chewing, ‘space’ for swallowing). After the tasting, they scored the perceived intensity of aroma. In this paper, we refer to the 0-120 s interval as a single tasting.

For the first session, 2mL of the diluted solution (0.4 g.L^-1^) were presented in a cup (30 mL) closed with a cap. The subject first kept the product in the mouth for 30 seconds without any action (0-30 s interval). During the next 30 seconds (30-60 s interval), the subject simulated chewing the aqueous solution by moving their jaws (this is noted here ‘jaw moving’ phase). This phase was set up in order to facilitate aroma delivery to the nasal cavity and the subject had to press ‘m’ at each jaw move. The subject then swallowed the product all at once and, depending on the FOP, either swallowed as frequently as possible (‘freq’ FOP) or as infrequently as possible (‘rare’ FOP) (60-120 s interval) without disrupting the subject. The solution presented in a cup was used to compare two swallowing behaviors — frequent versus infrequent swallowing — after ingestion, thereby allowing us to isolate the effect of swallowing frequency on post-deglutition aroma release.

For the second session, a gustometer (Multistimulator OG001) supplied by Burghart™ was used to deliver the solution in a controlled way into the subjects’ mouths via a PTFE tube. This gustometer consists of three taste modules, which are software-controlled syringe pump systems. It allows precise control of the flow rate, timing, and volume of solution delivered into the subject’s mouth, with a volume accuracy down to a few microliters and a temporal precision to less than 50 milliseconds. One module was used for delivering Evian® water, and the second module was used for delivering the aroma solution (0.4 g.L^-1^). The third module was not used. The subject placed the gustometer tubes on their tongue and gently stabilized them with their teeth, without biting down. The solution (2 mL) was then delivered at a constant rate for 1 minute (at a rate of 2mL.min^-1^). During the first minute (0-60 s interval), the subject either swallowed the product immediately as it entered the mouth (i.e., nearly continuously, “cont” FOP), or allowed the product to accumulate until the 60-second signal (imposed 60 s swallow) indicated that it should be swallowed (“noth” FOP). Then, the subject continued swallowing regularly until the end of the acquisition (e.g., every three breathing cycles, not strictly imposed). The solution delivered by gustometer was used to compare two swallowing behaviors — frequent versus infrequent swallowing — before final ingestion, thereby allowing us to isolate the effect of swallowing frequency during consumption (e.g., during the melting of a solid food).

For the third session, the subject kept the gelatin gummy disc in the mouth for 30 seconds without actions or movements, except for an imposed swallowing of saliva only, after 15 seconds (imposed 15 s swallow). This imposed swallow was set up in order to facilitate aroma delivery to the nasal cavity. During the following 30 seconds (30-60 s interval), the subject either sucked (“succ” FOP) or chewed (“chew” FOP) the product depending on the condition. The subject then swallowed the product and continued swallowing at a regular rhythm (e.g., every three breathing cycles, not strictly imposed) during 1 minute (60-120 s). The “gelatin gummy disc” delivery mode was used to compare two behaviors — chewing versus suction — before ingestion, thereby allowing us to isolate the effect of swallowing frequency during consumption.

Regarding the order of presentation of FOPs, the subject used a first food oral processing (‘chew’ for example, see Fig 1.b.) during the first 3 replicates, then a second one during the last three replicates (‘succ’ for example). The FOP conditions were randomized across subjects: for example, some participants performed ‘chew’ first and then ‘succ’, while others performed ‘succ’ first and then ‘chew’.

#### 2.1.5. Analysis of exhaled air

The exhaled air of the subjects was constantly analyzed using a PTR-ToF-MS device during all the tastings.

The PTR-ToF-MS (PTR-ToF-8000, Ionicon Analytik GmbH, Innsbruck, Austria) upgraded with the ion funnel to better focalize the ions into the ToF mass analyzer, ionized volatile compounds through proton transfer from H₃O⁺. The resulting ions were then analyzed by a time-of-flight mass spectrometer (ToF-MS). Parameters of the PTR-ToF-MS instrument were as follows: drift pressure of 2.3 mbar, drift temperature of 80°C, and drift voltage of 390 V, resulting in an electric field strength to number density ratio (E/N ratio) of 115 Townsend (Td, 1 Td = 10^−17^ V.cm^2^). Data were collected using the TofDAQ software provided by the manufacturer of the PTR-ToF-MS. Data acquisitions were performed at 1 mass spectrum ranging from m/z 0 to 227 per 100 ms (10 scans/s).

Nosespace sampling was performed using a custom-made lightweight headset equipped with two Teflon nozzles positioned at the entrance of the subject’s nostrils, directing exhaled air toward the PTR-ToF-MS transfer line. The system allowed subjects to move their heads freely and did not interfere with breathing or chewing. The PTR-MS transfer line and PEEK tubing of the helmet were maintained at 110 °C and 70 °C, respectively, and sampling was conducted at a total flow rate of 400 mL.min⁻¹. Connection to the PTR-MS was verified by monitoring naturally exhaled isoprene (m/z = 69.069877).

PTR-MS data was preprocessed with PTRMSR R package (code freely available on https://github.com/Chemosens/PTRMSR) based on a table of integration to extract the counts for several ions: isoprene, acetone, isoamyl acetate (ACI, m/z=131.107) and fragments (Appendix 1). The concentrations were estimated with a formula delivered by the PTR-MS manufacturer (Ionicon) and mainly consisted in a ratio with the concentration in H_3_O^+^. Each curve of aroma release was plotted to check the correct import of data.

#### 2.1.6. Declarative dynamic events and perceived final intensity

Subjects were instructed and monitored using PsychoPy software, which recorded swallowing events in real time and allowed subjects to rate aroma intensity at the end of each tasting. PsychoPy software (version 2.4. 2024) is an open-source Python-based software designed for building and running experiments in behavioral sciences. It allows precise control of stimuli presentation and response recording. In this study, PsychoPy was used to time instructions and collect subjects’ declarative inputs, specifically the timestamps of swallowing and mastication events during the tasting sessions.

Note that only the main product swallowing (60 s) and the 15 s one for the gummy disc were imposed; all other swallows were self-reported by the subjects.

A final exploratory question assessed perceived aroma intensity: ‘During this tasting, how intense did you perceive the aroma?’ Responses were recorded on a continuous scale ranging from 0 (not intense at all) to 10 (very intense). This question did not specify a particular moment during the tasting but rather referred to the overall perception.

### 2.2. Statistical analyses

Data were mainly analyzed with R software (R.4.4.1), but breath analysis and peak detection were conducted in Python with SciPy library. Data are shared on research.data.gouv.fr (Peltier, Caroline, 2026, “Dataset of aroma release *in vivo* (Digimouth)”, https://doi.org/10.57745/OVC3RL) and the R scripts are freely available on https://github.com/ChemoSens/ExternalCode/Digimouth.

#### 2.2.1. Analysis of aroma release curves

##### 2.2.1.1. Effect of FOP on AUC and Imax

For each delivery mode and each interval of time (0-30s, 30-60s and 60-120s), two indicators of aroma release were computed: the maximum intensity (Imax) and the area under the curve (AUC). Wilcoxon tests were conducted to identify differences between food oral processes.

##### 2.2.1.2. Release curves averaging

Release kinetics were analyzed. First, all curves were binned at 1-second intervals and then averaged within each FOP (n=3*30=90 curves, for 3 replicates and 30 subjects). The standard error of the mean (SD / √n) was computed and represented as a shaded ribbon around the mean curve.

#### 2.2.2. Perceived intensity analysis and relations with AUC

The effect of FOP on declared intensity was tested with the Wilcoxon test. The distribution of the results was presented as boxplots.

Then, relations between AUC and perceived intensities were studied with correlations.

First, Spearman correlations were calculated between AUC and intensities by delivery mode for all tastings to indicate whether, for a given delivery mode, a higher AUC indicates a higher perception.

Second, Spearman correlations between declarative intensity and AUC of aroma release curves were computed per subject (3 matrices * 2 FOP *3 replicates = 18 observations), then averaged over subjects. This calculation indicates whether, within a single subject, intensities were correlated to AUC.

Finally, this approach based on Spearman correlations was also conducted by delivery mode (6 observations per correlation coefficient).

#### 2.2.3. Clustering of subjects based on their AUC during the ‘jaw moving’ phase of ‘solution in a cup’ delivery model (30-60s)

Preliminary experiments and pilot tests revealed substantial inter-individual variability in aroma release during the “moving jaw” phase of the solution in a cup, likely related to differences in velopharyngeal opening. Based on these observations, a clustering approach was implemented to identify potential groups of subjects during this phase. Prior to analysis, the data were log-transformed using a base-10 logarithm, with a constant of +1 added systematically to all values to avoid negative inputs. The resulting time series were then smoothed by averaging the data within each one-second interval. Consequently, each time point represents, for each subject, the mean of the six repeated tastings of the same solution. From these averaged curves, the AUC was calculated for each subject, and a distance matrix was generated. Hierarchical clustering (Euclidean distance, Ward’s method) was then applied, revealing two distinct groups. These clusters were subsequently characterized using (i) the raw, non-preprocessed data and (ii) the log-transformed averaged data (iii) their intensity scores (iv) their frequency of ‘jaw moving’.

#### 2.2.4. Analysis of inter and intra-individual variability

##### 2.2.4.1. Analysis of breathing

Breath analysis was computed only on the 30 first seconds before putting the sample in the mouth (−30 - 0 s), so it was not influenced by food oral processing. Signal peaks were detected using the “scipy find.peaks” Python function for detecting breathing cycles.

Distances between two successive peaks were extracted then averaged for each tasting (on the -30-0s interval). These values were calculated for each subject, for all delivery modes and including warmup, then averaged. Inter- and intra-subject variability were quantified using the standard deviations across subjects (with average results by subject) and across repetitions (within each subject), respectively. The ratio SD_inter / SD_intra was used as a descriptive measure to characterize the relative contribution of between-subject variability. The reported ratios therefore provide a robust descriptive assessment of variability.

##### 2.2.4.2. Analysis of the variability of AUC and Imax of aroma release curves

Inter- and intra-subject variabilities were quantified using the standard deviations across subjects and across repetitions, respectively for AUC and Imax. The ratio SD_inter / SD_intra was also calculated.

Then, replicates were averaged and non-parametric tests were conducted (paired Wilcoxon tests) to assess the statistical differences between FOP. Boxplots were plotted to visualize the distributions of these parameters.

##### 2.2.4.3. Declarative behavior and relationship with aroma release

The declarative times of chewing and swallowing were recorded with PsychoPy software. First, the frequency of chewing and swallowing was computed for each interval of time (0-30s, 30-60s, 60-120s) in order to check if the subjects followed the instructions. A table documenting inter- and intra-individual variability was also computed as in Sections 2.2.3.1 and 2.2.3.2. Furthermore, the average number of additional reported swallows during the 0–60 s period was computed for each delivery mode.

#### 2.2.5. Missing data

Missing data occurred for two reasons: (i) subject B477 missed the gustometer session, and (ii) a technical issue occurred during the gelatin gummy disc acquisition for subject E809. Consequently, the final dataset consisted of 30 × 6 aroma release curves for the solution in a cup conditions, and 29 × 6 aroma release curves for the solution delivered by gustometer and gummy disc conditions (1 delivery mode, 2 FOP with 3 replicates = 6 tastings by session missing for the indicated subjects).

## 3. Results

### 3.1. Example of individual curves

Figure 2 shows the aroma release over 120 seconds for a single subject and a single replicate for each FOP and each delivery mode. For the solution under the “freq” protocol, during the 0-30 s phase (no oral processing), no aroma release is observed. In the 30-60 s phase (simulating mastication, with each chew indicated by a red striped line), a clear aroma release occurs. After swallowing the product (bold blue line), the subject performed seven swallows, indicated by blue dotted lines (Fig. 2a). Under the “rare” protocol (no swallowing), the first two steps are the same. Here, each declared chew corresponds clearly to a peak in aroma release. A small initial peak is observed at the very beginning of tasting (around 2 s), likely due to sniffing of the solution before placing it in the mouth (orthonasal pathway).

**Figure 2:**
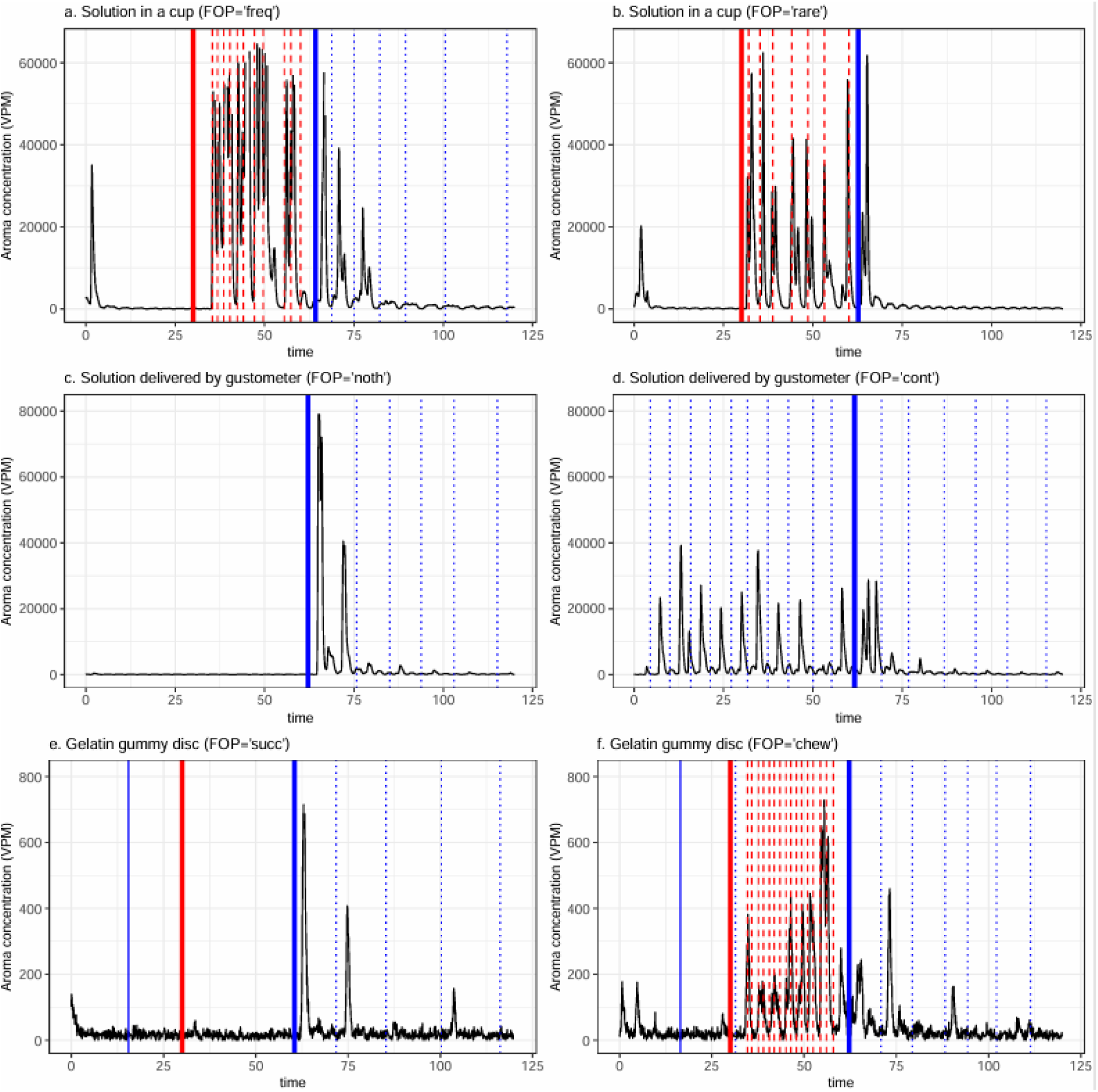
Example of aroma-release-over-time data for the first replicate of one subject (Q407) across the different FOP conditions: (a) solution in a cup with high-frequency swallowing (“freq”), (b) solution in a cup with low-frequency swallowing (“rare”), (c) solution delivered by gustometer, no swallowing (“noth”), (d) solution delivered by gustometer with continuous swallowing (“cont”), (e) gelatin gummy disc with suction (“succ”), and (f) gelatin gummy disc with a mastication protocol (“chew”). The red bold line indicates 30 seconds and the blue bold line indicates 60 seconds, corresponding to the times of product swallowing (solid or liquid). Red striped lines represent declared chewing movements, and blue dotted lines represent swallowing events. Blue continuous lines are imposed swallowing events.

For the gustometer, only two phases were considered: no oral processing (0-60 s), followed by swallowing (60-120 s, either all at once or continuously). During the first phase, no aroma is released under the “nothing” protocol, while in the “continuous swallowing” protocol, each swallowing event induces a peak in aroma release. In the second phase, the aroma signals are higher for the “nothing” protocol (Fig. 2c and 2d).

Finally, for the gelatin gummy disc, the aroma signal is overall much lower and noisier. Chewing appears to release aroma more effectively than suction (Fig. 2e and 2f).

### 3.2. Effect of FOP on AUC and Imax

Solutions in a cup (60–120 s) showed higher AUC values under the “freq” condition compared with the “rare” one (p<0.001) (Figure 3a). Most subjects displayed a consistent within-subject increase of AUC when swallowing more frequently, indicating that swallowing events strongly drive aroma clearance and subsequent release.

**Figure 3:**
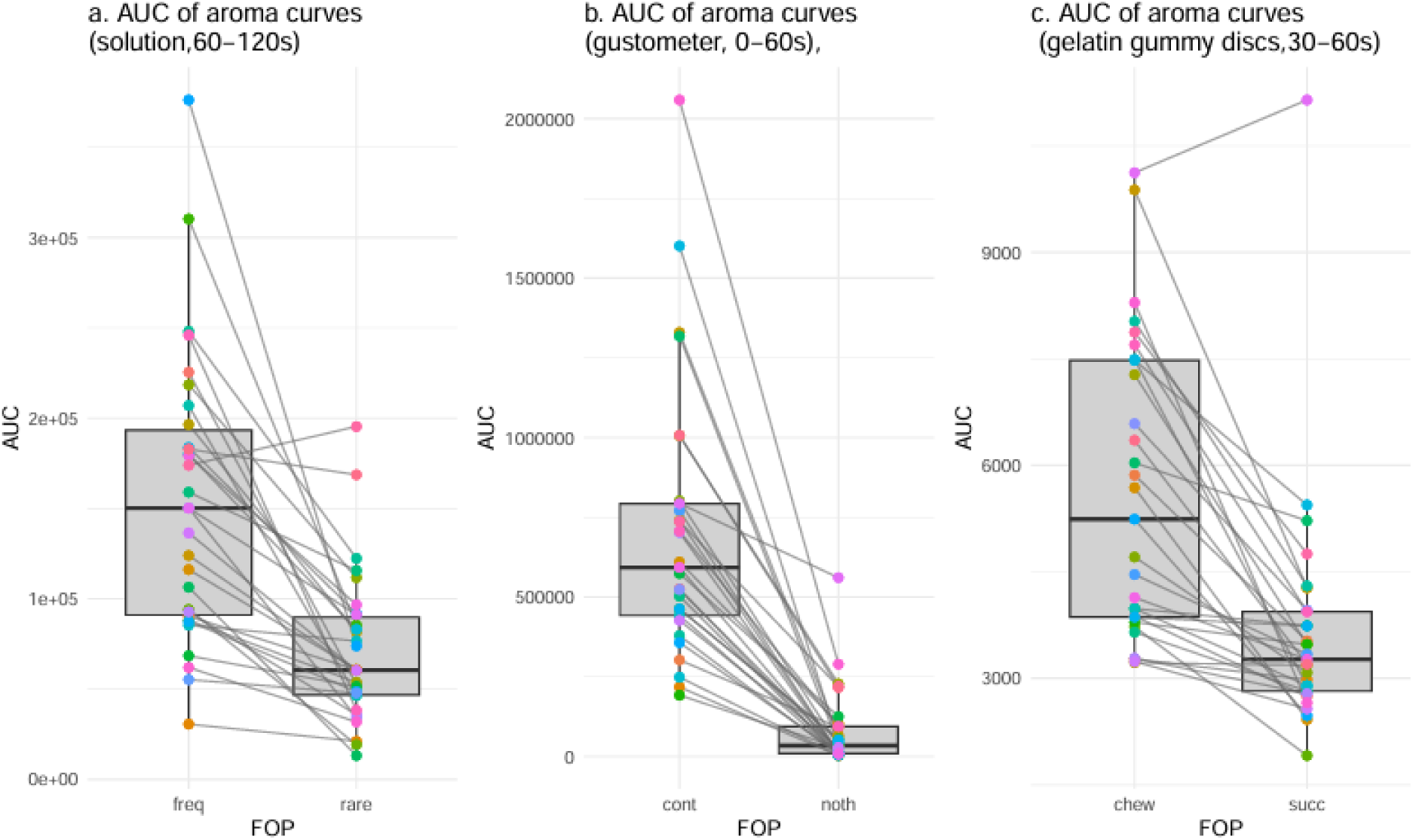
Comparison of averaged AUC values from aroma release curves under different conditions (replicates were averaged). (a) averaged AUC values for ‘solution in a cup’ measurements between 60–120 seconds, comparing frequent (“freq”) and “rare” FOP events. (b) averaged AUC values for ‘solution delivered by gustometer’ measurements between 0–60 seconds, comparing continuous (“cont”) and no-triggered (“noth”) swallowing events. (c) averaged AUC values for gelatin gummy disc measurements between 30–60 seconds, comparing chewing (‘chew’) and suction (“succ”) FOP. Each colored dot represents a subject, and lines connect paired observations from the same subject.

For the solutions delivered by gustometer (0–60 s), the continuous swallowing “cont*”* FOP produced markedly higher AUCs than the “noth” condition. The contrast was especially pronounced, with very low AUC values when subjects did not swallow (p<0.001). This confirms that, in the absence of swallowing, the transfer of volatile aroma compounds from the mouth into the nasal cavity is strongly limited (Figure 3b).

For gelatin gummy disc (30–60 s), AUC was higher in the “*chew”* condition than in *“succ”*. The difference was smaller than for solutions in a cup or delivered by the gustometer, but highly significant for most subjects (p<0.001). Chewing generated stronger mechanical breakdown and greater aroma release compared with suction (Figure 3c).

Overall, the paired lines indicate that most subjects exhibited the same directional effect, even though the magnitude differed widely. This reinforces the idea that the FOP imposes a strong mechanical control over aroma release, but with substantial inter-individual variability.

### 3.3. Release curve averaging

The averaged aroma release curves (Fig. 4) indicate that the main observations obtained for a single subject and replicate in Fig. 2 remain valid across the studied population, after individual variations due to breathing and swallowing are smoothed out.

**Figure 4:**
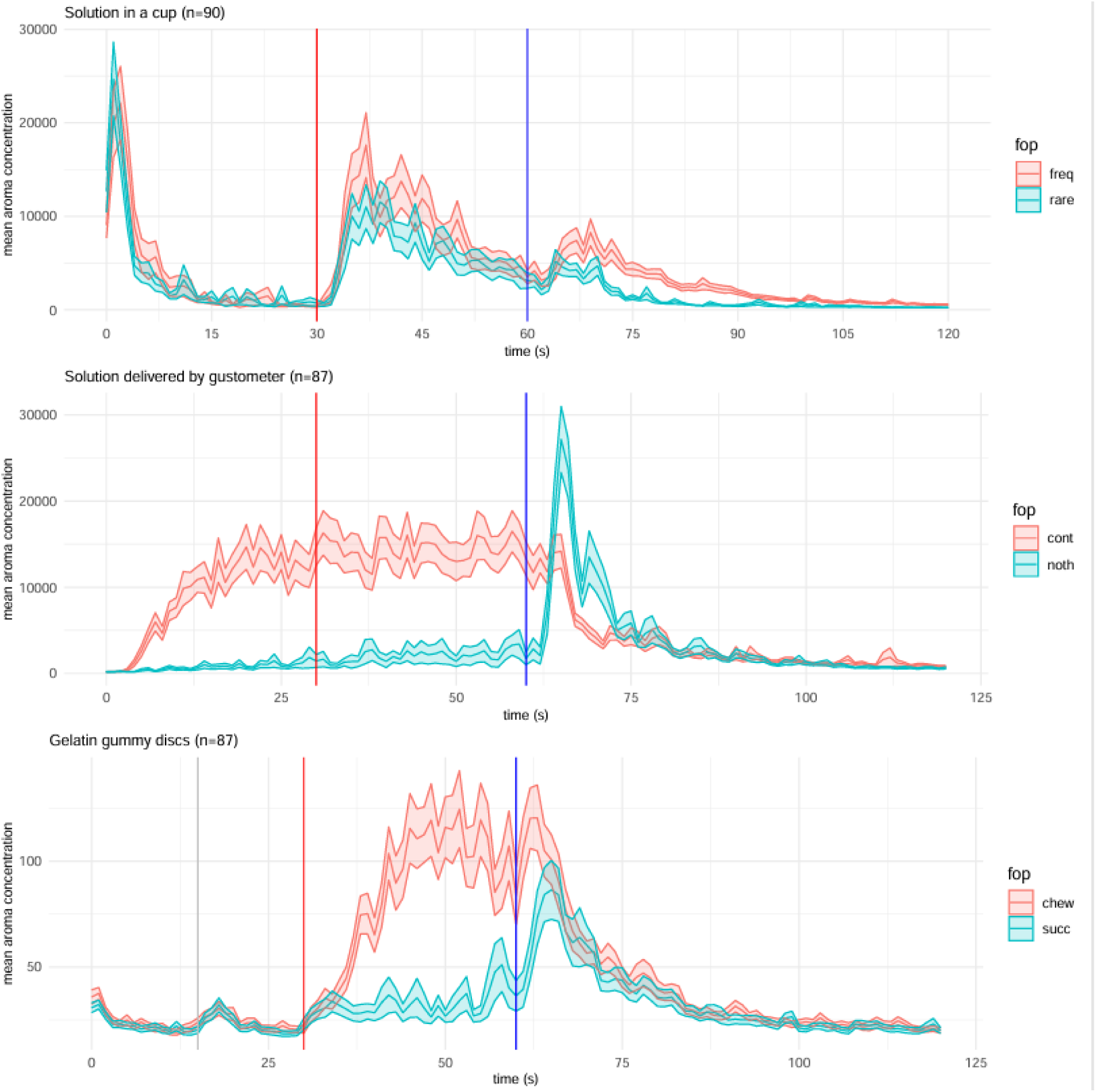
curve of averaged aroma release for all tastings with standard errors of the mean. The red bold line indicates 30 seconds and the blue bold line indicates 60 seconds, corresponding to the times of imposed product swallowing.

For the solution in a cup, an initial aroma peak was observed during the 0-30s interval: this is probably due to the fact that the participants sniffed the aroma compounds (orthonasal pathway) from the samples by natural breathing before ingestion, and this was consequently detected by the PTR-MS. Aroma release then increased when mimicking chewing (30-60 s). Significant differences were found between the ‘freq’ and ‘rare’ swallowing protocols: aroma release was found to be higher and lasted longer when swallowing occurred frequently in the 60-120 s interval (p<1·10^-9^).

For the solution delivered by gustometer, significant differences were observed between the continuous swallowing (‘cont’) and the no action (‘noth’) protocols. Under continuous swallowing, the aroma release remained relatively constant throughout the tasting period (0-60 seconds) and decreased only after the last swallowing (60 s). In the no action protocol, little aroma was released before swallowing, followed by a sharp peak once swallowing occurred.

Finally, for the gelatin gummy disc, chewing clearly released more aroma than suction, highlighting the strong influence of mastication on aroma release.

### 3.4. Perceived intensity analysis and relations with AUC

For any given delivery mode, the intensity scores (0–10 scale) provided by the subjects did not differ significantly between FOPs (Fig. 5).

**Figure 5:**
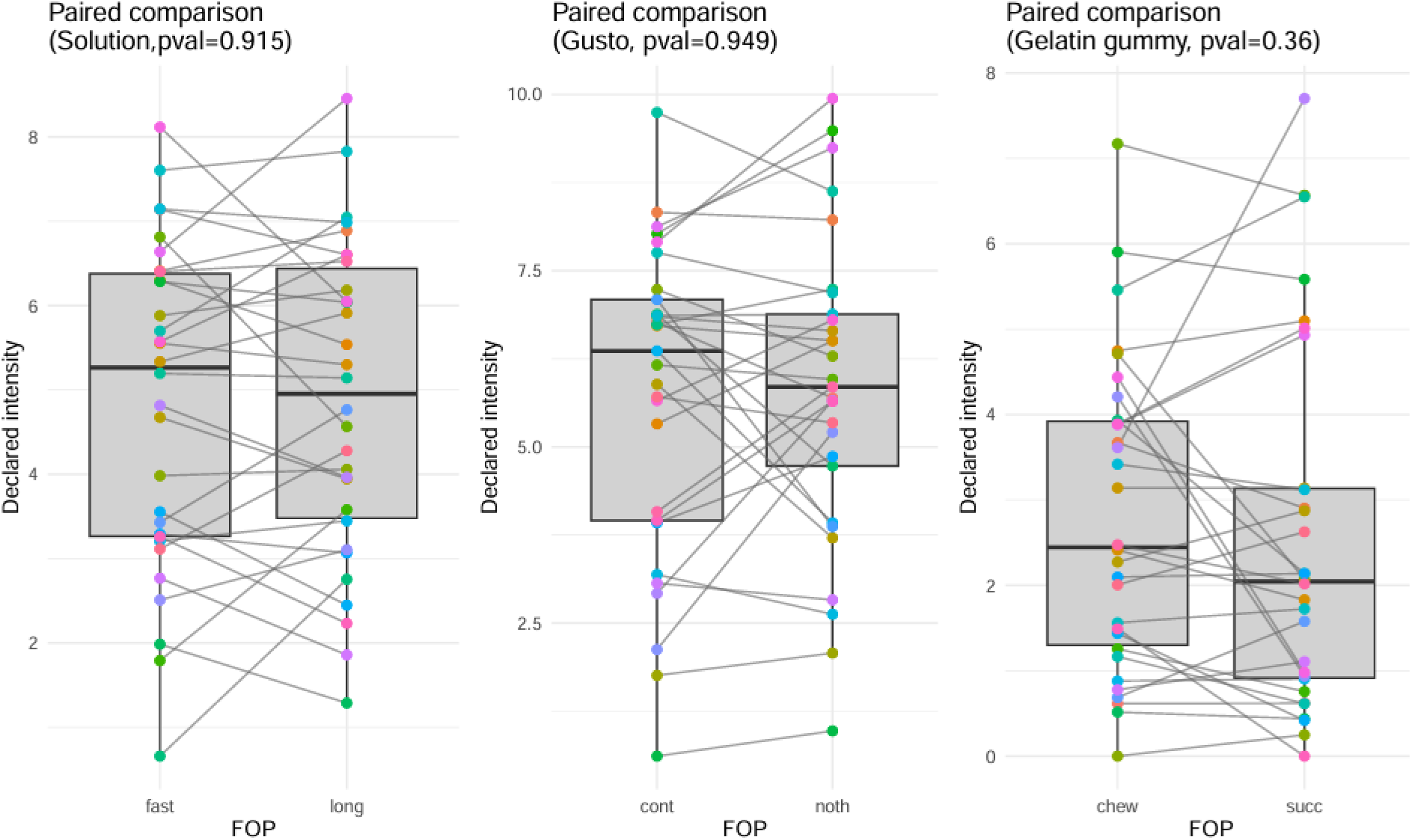
Distribution of declared intensity by delivery mode and FOP.

The panel correlation between AUC and intensity was 0.4 indicating that there is a positive correlation between intensity and AUC. However, looking at delivery mode level, the correlation coefficients were drastically weaker (0.09 in average for the solution with a cup, - 0.04 for the solution delivered by the gustometer and -0.02 for the gelatin gummy disc).

The average Spearman correlation between the 18 measured AUC and the 18 declared intensities across subjects was 0.53, indicating that within each subject, perceived intensity and AUC values were positively correlated.

When the correlation was computed separately for each delivery mode, the values dropped to 0.12 for solution in a cup, -0.009 for solution delivered by gustometer, and 0.10 for the gelatin gummy disc.

This suggested that the overall correlation observed across subjects was mainly driven by the large differences in both intensity scores and AUC for the gelatin gummy disc. In other words, the correlation appeared to be strong only when the delivery modes differed markedly, while considering only the variability of aroma release inside a given delivery mode did not lead to strong correlations.

### 3.5. Clustering of subjects based on their AUC during ‘jaw moving’ phase of the solution in a cup tasting (30-60s)

During the ‘jaw moving’ phase, subjects held the solution in their mouths without swallowing, while moving their jaws as if chewing a solid product. Hierarchical clustering identified two groups clearly displaying distinct behaviors: Group 1 (G1, n = 22) and Group 2 (G2, n = 8).

These groups are illustrated in Figure 6, which highlights their divergent response patterns. Figures 6a and 6b show the raw release curve of a subject in G1 and another one in G2, showing a clear difference between these subjects (the two figures have the same scale), and showing negligible signal for the subject in G2 compared to G1 in the 30-60s interval. This observation of the raw data was unexpected, but justified the cluster analysis previously presented. In Figure 6c, the averaged intensity profiles are shown with their corresponding standard deviations and showing that Group 1 was characterized by subjects exhibiting a strong increase in signal intensity during the “jaw moving” phase. This response suggested that these subjects may open the velopharyngeal port during this phase. In contrast, Group 2 displays very small signal during this phase, indicating that these subjects likely maintain better velopharyngeal closure.

**Figure 6:**
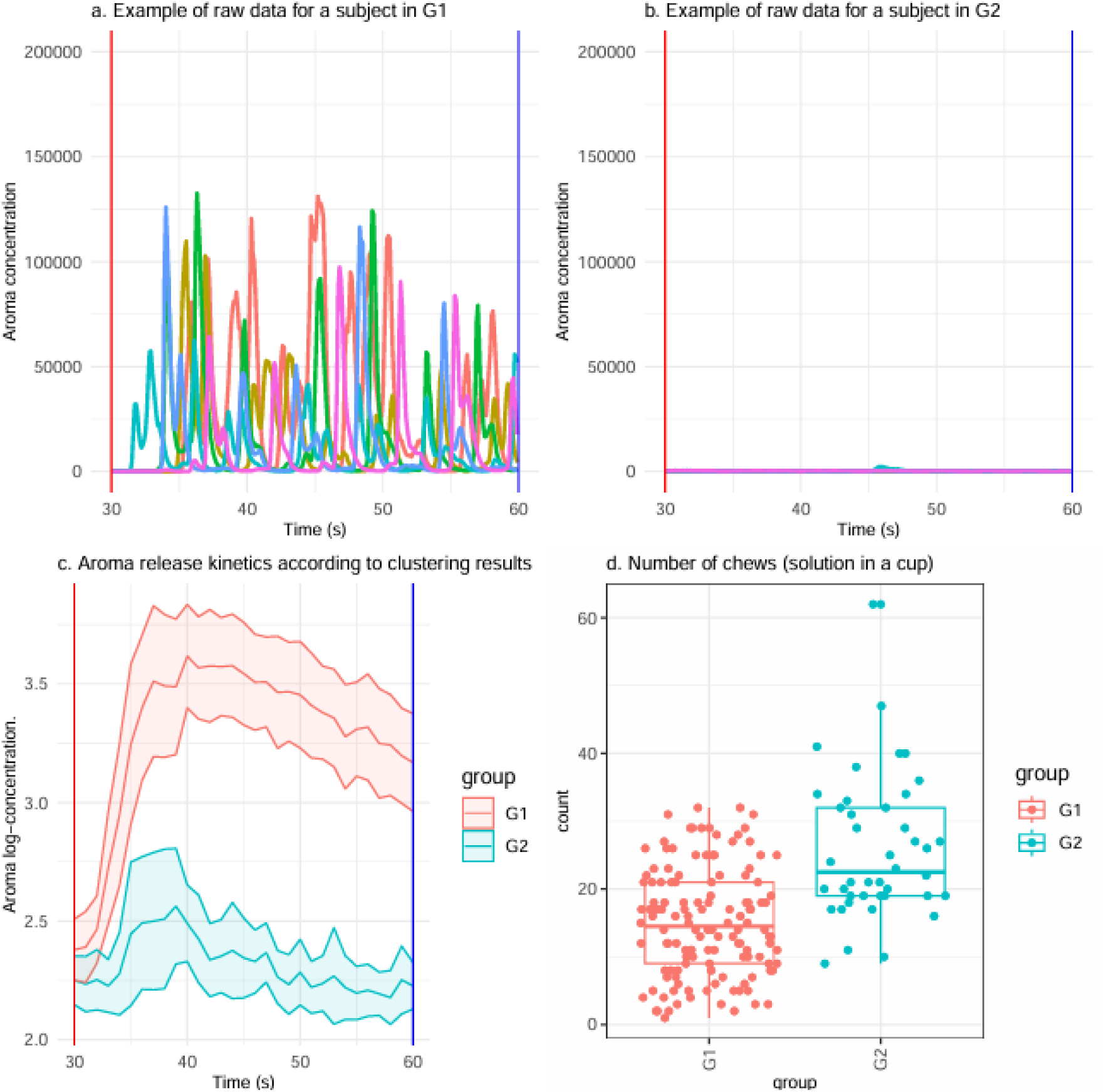
Comparison of aroma release and chewing behavior between groups G1 and G2 for the solution in a cup, with the two FOPs. (a) Raw concentration data for a subject from G1 (Q407) during chewing period (30-60) in solution (b) Raw data for a G2 subject (T386) (c) Average aroma release kinetics (log[1+concentration]) for both groups, with mean lines, 95% confidence intervals, and key chewing cycle times (starting chewing moves in red, swallowing in blue) (d) Number of chews recorded per subject in both groups, shown as individual points and summarized with boxplots (median, quartiles, range).

Statistically significant differences were observed (Mann–Whitney U test) between the number of chews for G1 and G2 (p<0.001): surprisingly, G2 was “chewing” significantly more frequently than G1 (Fig. 6d). No statistical difference was observed in the perceived intensity.

### 3.7. Analysis of inter and intra-individual variability

#### 3.7.1. Analysis of breathing

During the -30 to 0 s interval, an average breathing cycle of 3.33 s (SD = 1.7; range 1–9), corresponding to 18 peaks per minute was observed.

#### 3.7.2. Analysis of the variability of AUC and Imax of aroma release curves

Across all matrices and FOPs, inter-subject variability was consistently higher than intra-subject variability for both AUC and Imax, with ratios generally ranging from 2 to 4 (Table 1). Supplementary values (mean, standard deviations) are available in Appendix 2. For AUC, solution in a cup and gelatin gummy disc samples showed ratios around 3, while the continuous-swallow gustometer (“cont” FOP) displayed the highest value (4.39). For Imax, ratios were slightly lower overall (≈1.5–3.17), but the pattern remained similar: inter-subject differences dominated the total variability regardless of the delivery mode and the FOP. AUC captured the overall aroma release over time by including both intensity and duration. It might have been more sensitive to differences in swallowing behavior, bolus residence time, and individual kinetics. This often resulted in higher inter-individual variability and a broader dynamic range.

**Table 1:** Ratios of inter-/intra-subject standard deviation, quantifying the relative contribution of between-subject variability compared to replicate tastings by the same subject (see Appendix 2 for averages and standard deviations).

|  | Delivery mode | FOP protocol | Ratio |
| --- | --- | --- | --- |
| AUC | Solution in a cup | rare | 3.52 |
|  |  | freq | 3.01 |
|  | Solution delivered by gustometer | noth | 2.69 |
|  |  | cont | 4.39 |
|  | Gelatin gummy disc | chew | 2.80 |
|  |  | succ | 3.75 |
| I<sub>max</sub> | Solution in a cup | Rare | 1.94 |
|  |  | Freq | 1.72 |
|  | Solution delivered by gustometer | Noth | 1.54 |
|  |  | Cont | 3.17 |
|  | Gelatin gummy disc | Chew | 1.72 |
|  |  | Succ | 1.79 |

Imax reflects only the peak intensity of release. It is less influenced by the temporal shape of the curve and therefore tends to produce lower ratios. In this study, AUC was more discriminant across FOPs (especially for continuous swallowing).

#### 3.7.3. Analysis of declared chews and swallows and relations with aroma release

Table 2 shows that intra-subject variability is much lower (more than 3 times) than inter-subject variability for all declarative indicators, except in the “rare” FOP. In this case, this can be explained by the fact that subjects swallow mainly only once when consuming the product. Consequently, only little variability was observed and probably due to occasional ‘accidental swallows’ that could occur potentially in any subject. Furthermore, Table 2 shows that the subjects followed the instructions: the average frequency of swallowing in the “frequent” condition is 10 times higher than in the “rare” condition. The “regular” condition was intermediate (5 or 6 events per minute).

**Table 2:** Mean values and inter/intra-subject SD ratio, quantifying the relative contribution of inter-subject variability compared to replicate tastings by the same subject.

| Declared event | Delivery mode | FOP<br>protocol | Mean | ratio |
| --- | --- | --- | --- | --- |
| chew (30-60s) | Solution in a cup | 'chew' | 17 | **4,75** |
|  | Gelatin gummy disc | 'chew' | 16,5 | **5,6** |
| Swallowing (60s-120 s) | Solution in a cup | 'rare' | 1.71 | **1,88** |
|  | Solution in a cup | 'freq' | 9.9 | **3,9** |
|  | Solution delivered by<br>gustometer | 'noth' | 5.97 | **5** |
|  | Solution delivered by<br>gustometer | 'cont' | 6.34 | **3,15** |
|  | Gelatin gummy disc | 'chew' | 6.09 | **4,92** |
|  | Gelatin gummy disc | 'succ' | 5.64 | **5,47** |

As an additional indicator, during the 0–60 s phase (when no swallowing was requested for the solution in a cup and gelatin gummy disc), the number of swallows for the solution (including the 6 tastings) was as follows: 0 for 19 subjects, between 1 and 5 for 5 subjects, 6 to 8 for 4 subjects, and more than 10 for 2 subjects. For the gelatin gummy disc, a similar pattern was observed: 21 subjects reported no additional swallowing, while 2 subjects reported more than 10 swallowings events accumulated across the six tastings. For the solution delivered by gustometer, during the ‘no action’ protocol (“noth”), 20 subjects reported 0 swallows, and the remaining subjects reported between 1 and 4. Finally, under the continuous swallowing condition (“cont”) in the gustometer, the average number of swallows per tasting was 11 (range 5–24). This indicated that the subjects followed the instructions correctly.

## 4. Discussion

### 4.1. Overview of the main results and interpretation of the results

First, food oral processing (FOP) substantially affected aroma release for all delivery modes. In particular, swallowing was the main contributor to sharp peaks in VOC concentration, in line with studies showing that deglutition generates bursts of retronasal airflow and promotes aroma transport toward the nasal cavity (Buettner et al., 2002; Tarrega et al., 2011). Chewing also increased release, likely due to product breakdown and enhanced mass transfer or also circulating the flavored air toward the nose, for subjects who do not achieve proper velopharyngeal closure (Buettner, 2001; Buettner et al., 2002; Tarrega et al., 2011).

Second, as commonly reported, linking aroma release metrics such as AUC or release kinetics to perceived intensity remains challenging. The very low correlation coefficients calculated for each subject and delivery mode illustrated this difficulty. Several factors may account for this weak association. First, the panelists were not trained and may therefore have used the intensity scale inconsistently. Second, these analyses were performed on a single delivery mode, under the assumption that, for this given delivery mode, greater aroma release leads to higher perceived intensity. However, when differences in aroma release are too small, they may fall within the range of experimental noise or below differential sensory thresholds rather than reflect meaningful sensory differences, making such comparisons of limited relevance. Finally, the way in which individuals cognitively integrate the sensory signal remains unknown: judgments may rely on the total area under the curve, the maximum concentration peak, or other temporal or qualitative cues likely different from one individual to another. Sensory integration is thus likely more complex than what can be captured by a single AUC-based descriptor and warrants further investigation.

The presence of two distinct groups during the ‘chewing’ phase for solution illustrates that some people do not close entirely their velopharynx during these periods whereas others do, as presented by Repoux et al. (2012). It seems that those with good velopharyngeal closure made chew movements much faster than the others. This observation has not been reported in the literature and may warrant further investigation. For example, does mimicking faster chewing lead to velopharyngeal closure in subjects?

The suction protocol results in weaker aroma release than the chewing protocol. This may be consistent with the fact that the exchange area between the product and the air increases more during chewing than during suction. Indeed, aroma release is theoretically related to this exchange area. Another possible explanation is that the velopharyngeal opening is greater during chewing than during suction (at least for some subjects).

The breathing cycles observed in this study are slightly faster than the ones presented in the literature. Sembuligam and Sembuligam (2012) indicate a respiratory rate of 12-16 cycles/minute and our results show a respiratory rate of 18 (60/3.33s). This could be due to the fact that the subjects were stressed by the connection to the PTR-MS and following the instructions. Furthermore, the rhythm is evaluated on a 30 s period that is short for evaluating a frequency of cycle by minute.

### 4.2. Limitations of the study

Here, we conducted our study only on isoamyl acetate (m/z = 131.107), however, a preliminary analysis showed that other ions were present in the signal, namely m/z = 71.08 and m/z = 61.028, which are fragments of isoamyl acetate. Possible improvement could consist in a co-analysis of isoamyl acetate and its fragments.

The aroma concentration in the gelatin gummy discs was not strictly standardized due to possible losses during disc preparation and should be improved in future experiments. Further experimentations could consider the real concentration in the gelatin gummy discs after manufacturing.

Oral events were self-reported, which introduces temporal uncertainty. As the swallowing can occur only before an expiration, swallowing times could potentially be refined by aligning them with breathing cycles. This could be useful for modeling aroma release. Further works could also include using motion capture instrument (Optotrak type) or filming the panelists to re-align the reporting of times or to reduce their cognitive load and ask to concentrate on other tasks such as reporting perceived intensity versus time.

Panelists were not trained for intensity evaluation, and no dynamic sensory data were collected. Time–intensity procedures were tested in preliminary work but abandoned due to task complexity; we prioritized collecting reliable swallowing and chewing annotations. Still, it would be interesting to reproduce such experiments with gustometer’s concentrations changing in time and acquiring time-intensity signals, while using sensors to automatically detect chews and swallows and thus reduce the cognitive load of the panelists. With a simple protocol such as continuous swallowing, the time-intensity could be recorded, for example for increasing concentrations. Other protocols such as discontinuous time-intensity methods could also be considered.

### 4.3. Perspectives

As product composition, including sugar and acid content, is known to modulate release kinetics and perception, further work could investigate the impact of adding more aroma or sapid compounds in the products. Additional physiological measurements (oral and nasal cavity volumes, airflow, salivary flow, ethnicity) would help generalize these findings and characterize the different groups observed.

One of the objectives of this study was to produce data usable for mechanistic modeling to bridge *in vitro* and *in vivo* predictions of individual aroma-release curves. Mechanistic modeling is essential to translate experimental aroma release data into predictive tools capable of describing aroma release across food matrices and oral processing conditions. Such models require well-characterized experimental datasets that capture both the dynamics of aroma release and the associated sources of variability, particularly those related to oral processing and respiratory behavior. These data will be analyzed with these models. Incorporating velopharyngeal opening and inter-individual heterogeneity will be essential for accurate personalized predictions and will be the object of further works.

## 4. Conclusions

The inter- and intra-individual variability in aroma release was documented on three different matrices (aqueous solutions delivered in cups, with a gustometer and gelatin gummy discs) showing that inter-individual differences are higher than intra individual differences. The effect of swallowing was shown to be crucial for explaining aroma release kinetics. Highly significant differences were observed between protocols involving rare versus frequent swallowing. Moreover, aroma release was greater when the product was chewed than when it was only sucked. No aroma release was detected when no oral activity occurred, indicating that the velopharyngeal port remains closed during this period. Two groups of subjects were identified: those who open their velopharynx during jaw movement and those who keep it closed. These freely available data will be valuable for future mechanistic modelling.

## Acknowledgements

The research leading to these results has received funding under grant (24.7 DIGIMOUTH) from Carnot Qualiment© (DOI : 10.17180/h5gd-gk88) supported by Agence Nationale de la Recherche (20 CARN 0026). This work was performed thanks to CALIS network’s resources (doi: 10.15454/9PGN-W156). The authors thank Michel Tavan for technical help, and the panelists for their time and involvement.

**Appendix 1:** table of integration of aroma ions. m/z: mass-to-charge ratio of the volatile component, low: lower bound of integration of the peak, up: upper bound of integration of the peak

|  | [MH]<sup>+</sup> |  |  |
| --- | --- | --- | --- |
| name | m/z | low | up |
| Isoprene (2-methylbuta-1,3-diène) | 69.070 | 69.03429 | 69.11106 |
| Isoamyl Acetate (3-methylbutyl acetate) | 131.107 | 131.015 | 131.229 |
| Isoamyl Alcohol (3-methylbutan-1-ol) | 89.096 | 88.896 | 89.167 |
| C<sub>5</sub>H<sub>11</sub><sup>+</sup> (3 isomers of methylbutene, protonated) | 71.086 | 71.032 | 71.136 |
| C<sub>2</sub>H<sub>5</sub>O<sub>2</sub><sup>+</sup> (acetic acid protonated) | 61.028 | 60.984 | 61.082 |
| H<sub>3</sub>O<sup>+</sup> (hydronium ion) | 21.022 | 20.998 | 21.046 |
| H<sub>3</sub>O<sup>+</sup>(H<sub>2</sub>O) (water dimer protonated) | 39.033 | 38.988 | 39.07 |
| Acetone (propan-2-one) | 59.041 | 59.02949 | 59.06 |

**Appendix 2:** Mean and standard deviations for AUC and intensity during the whole period (0-120s)

|  | **Delivery mode** | **FOP protocol** | **Mean** | **Sd_total** | **Sd_intersubject** | **Sd_intrasubject** | **Ratio** |
| --- | --- | --- | --- | --- | --- | --- | --- |
| AUC | Solution in a cup | rare | 318186 | 250866 | 234197 | 66479 | 3.52 |
|  |  | freq | 481928 | 323291 | 296845 | 98776 | 3.01 |
|  | Solution delivered by gustometer | noth | 320324 | 244903 | 219328 | 81468 | 2.69 |
|  |  | cont | 905364 | 514823 | 493562 | 112434 | 4.39 |
|  | Gummy gel discs | chew | 5625 | 2265 | 2046 | 731 | 2.80 |
|  |  | succ | 3664 | 1727 | 1630 | 435 | 3.75 |
| Imax | Solution in a cup | rare | 79410 | 42287 | 36479 | 18843 | 1.94 |
|  |  | freq | 95429 | 44351 | 36654 | 21290 | 1.72 |
|  | Solution<br>delivered by<br>gustometer | noth | 97733 | 49699 | 39565 | 25668 | 1.54 |
|  |  | cont | 90379 | 42719 | 39835 | 12575 | 3.17 |
|  | Gummy gel<br>discs | chew | 566 | 274 | 224 | 130 | 1.72 |
|  |  | succ | 394 | 282 | 224 | 125 | 1.79 |

## Notes

https://doi.org/10.57745/OVC3RL

### Competing Interest Statement

The authors have declared no competing interest.

